# CXCR4 intracellular protein regulates drug resistance and tumorigenic potential via modulating the expression of Death Receptor 5

**DOI:** 10.1101/843995

**Authors:** Mushtaq Ahmed Nengroo, Shrankhla Maheshwari, Akhilesh Singh, Anup Kumar Singh, Rakesh Kumar Arya, Priyank Chaturvedi, Ayushi Verma, Krishan Kumar Saini, Annapurna Gupta, Anjali Mishra, Dipak Datta

## Abstract

CXCR4 overexpression in solid tumors has been strongly associated with poor prognosis and adverse clinical outcome. However, CXCR4 signaling inhibitor drug Plerixafor has shown limited clinical success in cancer treatment. Therefore, CXCR4 signaling may not be the exclusive contributor to its pro-tumorigenic functions. In our continuous effort to understand the chemokine receptor signaling inhibition as cancer therapy, here we unexpectedly discovered that instead of its signaling, intracellular CXCR4 protein augments therapy resistance and pro-tumorigenic functions. Unbiased proteome profiler apoptosis array followed by immunoblot, FACS, real-time PCR and ChIP analyses demonstrate that CXCR4 promotes DR5 downregulation via modulating differential recruitment of transcription factors p53 and YY1 to its promoter. Surprisingly, inhibiting CXCR4 mediated signals failed to block the above phenotype. Irrespective of CXCR4 surface expression, its loss compromised colon tumor growth *in vivo*. Finally, TCGA data mining and human patient sample data analysis showed CXCR4 and DR5 are inversely regulated in human cancers. Together, we showed evidence for the first time that targeting CXCR4 intracellular protein may be critical to dampen the pro-tumorigenic functions of CXCR4.

## Introduction

Cancer is the 2nd leading cause of death globally and was responsible for an estimated 9.6 million deaths in 2018 [1]. Drug resistance or inadequate chemotherapy response in patients is one of the pivotal reasons for cancer-associated colossal mortality and morbidity. Therefore, there is a dire need to understand the molecular mechanisms lying behind cancer therapy resistance. Cancer cells become therapy resistant by smartly evading the execution of apoptosis posed by chemotherapeutic drugs [2, 3]. The evasion of apoptosis by cancer cells largely relies on tilting the balance towards the increased expression of anti-apoptotic genes against pro-apoptotic genes [4, 5]. Death Receptors (DR) are critical pro-apoptotic factors present in healthy cells, immensely regulate cell death, and extensive literature suggests that paucity of death receptors in cancer cells promotes therapy resistance in diverse solid tumors [6-8]. So, the restoration of the functional death receptors is an attractive strategy to sensitize cancer cells against chemotherapy.

Over the last decade or so, chemokine receptor CXCR4 has been extensively reported to be overexpressed in most human solid tumors, and its association is strongly correlated with poor prognosis and adverse clinical outcomes [9, 10]. CXCR4 ligand (CXCL12) mediated signals are well established to explain its enormous role in modulating organ-specific metastasis of solid tumors [11-13]. Still, its effect on other pro-tumorigenic functions like therapy resistance and tumor-initiating capabilities are poorly understood. Moreover, multiple inhibitors of the CXCL12-CXCR4 axis, such as AMD3100 or plerixafor, and Nox-A12 have shown limited clinical success in cancer treatment [14]. These observations pose a serious concern regarding the contribution of the CXCL12-CXCR4 signaling axis in the pro-tumorigenic role of CXCR4. Earlier, we and others have extensively reviewed the dogma and opined that CXCR7, another CXCL12 high-affinity binding receptor could be the reason for the limited success of CXCR4 blockade in clinics [14-18]. Unfortunately, recent preclinical data for blocking both the chemokine receptors also failed to explain the pro-tumorigenic functions of these axes. In our relentless effort to understand the chemokine receptor signaling inhibition in cancer therapy [15, 19-22], here we surprisingly discovered that neither CXCR7 nor CXCR4-CXCL12 signaling axis is responsible for therapy resistance and tumorigenic potential, rather CXCR4 protein in the cancer cells plays a very critical role in positively modulating pro-tumorigenic functions such as therapy resistance across multiple solid tumors. Our intricate *in vitro* and *in vivo* experiments in multiple cancer settings for CXCR4 gain and loss of function studies demonstrate that CXCR4 protein but not its CXCL12 mediated signals modulate chemotherapy resistance and tumorigenic potential via inversely regulating the expression of DR5. CXCR4-DR5 inverse regulation was also validated in diverse human cell lines and human cancer patient samples, suggesting its possible therapeutic implications.

## Material and Methods

### Reagents and Antibodies

Doxorubicin, Paclitaxel, 5-fluorouracil, AMD3100, DAPI, anti-β-Actin (cat# A3854) antibody, Crystal violet dye, Lipophilic DiL, and DiO dyes, and Polybrene were obtained from Sigma Aldrich. Cisplatin was purchased from CADILA pharmaceuticals. ImmEdge pen (hydrophobic barrier pen), Bloxall blocking solution, were purchased from Vector Laboratories, Inc. Burlingame. Fluorochrome conjugated secondary antibodies, Accutase, ProLong™ Gold Antifade Mountant, and Verso cDNA synthesis kit, as well as SYBR® Green Real-Time PCR Master Mix, was purchased from Molecular Probes-Invitrogen. Taqman probes were purchased from Thermo Fischer scientific. DR5 (cat# 8074), YY1 (cat# 2185), p53(cat# 2527), Sp-1 (cat# 9389), Cleaved Caspase-8 (cat# 9496), p-ERK (cat# 9101) and GAPDH (cat# 2118) antibodies were procured from Cell Signaling Technology, Inc. CXCR4 antibody for western blotting was purchased from Abcam (cat# ab124824). HRP-conjugated secondary antibodies were obtained from Santa Cruz Biotechnology. APC conjugated CXCR4, CXCR7, PE-conjugated DR5, and IgG antibodies were purchased from BD Biosciences. Nonconjugated CXCR4, CXCR7 monoclonal antibodies, chemokine CXCL12, and CXCR4 antagonist peptide AMD3100 were procured from R&D systems. All chemicals and antibodies were obtained from Sigma unless specified otherwise.

### Cell Culture

Human breast cancer cell lines MCF7, MDA-MB-468, colon cancer cell lines DLD1, HCT 116, HT 29, COLO 205, SW 620, lung cancer cell lines A549, NCI-H-358 and prostate cancer cell line PC3 were obtained from American Type Culture Collection (ATCC), USA. Mycoplasma free early passage cells were resuscitated from liquid nitrogen vapor stocks and inspected microscopically for stable phenotype before use. MCF7, MDA-MB-468, DLD1, HT 29, COLO 2O5, A549, NCI-H-358, and PC3 cells were cultured in RPMI 1640 medium containing 10% fetal bovine serum (Gibco/Invitrogen), supplemented with anti-anti (Invitrogen). SW 620 cells were cultured in DMEM medium containing 10% fetal bovine serum (Gibco/Invitrogen), supplemented with anti-anti (Invitrogen). The HCT116 cell line was cultured in McCoy’s medium containing 10% fetal bovine serum (Gibco/Invitrogen), supplemented with anti-anti (Invitrogen).

### Cytotoxicity assay (SRB assay)

*In vitro* cytotoxic activities of Doxorubicin, Paclitaxel, 5-Fluorouracil, Cisplatin, TRAIL, and AMD3100 were assessed by using standard SRB assay as described before [23, 24]. The absorbance of the treated and untreated cells was measured on a multi-well scanning spectrophotometer (Epoch Microplate Reader, Biotek, USA) at a wavelength of 510 nm. Percent inhibition in cell growth was calculated by using the formula [100-(Absorbance of compound treated cells/Absorbance of untreated cells)] X 100.

### Flow cytometry

Cell surface expression of CXCR4, CXCR7 and DR5 in different cell lines was analyzed by Flow cytometry. In brief, cells were allowed to grow up to 70-80% confluence and then harvested with TrypLE (Invitrogen) for single-cell suspension in FACS buffer (PBS with 0.1% BSA). Cells were stained with fluorochrome-conjugated antibodies in FACS buffer for 30-45 min at room temperature in the dark. After washing and centrifugation, cell pellets were resuspended in FACS buffer and analyzed by FACS Calibur (BD). Acquired data were analyzed using FlowJo software (Treestar). Cell sorting was carried out by FACS Aria (BD).

### DiL DiO Staining

For staining the lipophilic dye DiL (1,1’-Dioctadecyl-3,3,3’,3’ Tetramethylindocarbocyanine Perchlorate) and DiO (3,3’-Dioctadecyloxacarbocyanine Perchlorate), cells were stained with 10μM of DiL (Red), and DiO (Green) dyes solution respectively for 45 min at 37°C. After that, cells were spun down, rinsed, and resuspended in fresh medium and seeded in 12-well plates. The next day, cells were treated with the chemotherapeutic agents at the desired concentration and visualized under the microscope at different timings or harvested for the analysis in FACS Calibur. For FACS analysis, DiL and DiO stained cells were harvested, washed, centrifuged, and resuspended in FACS buffer for analysis. Cells were analyzed under FL3 and FL1 channels in FACS Calibur.

### Western blotting

Cells were lysed in RIPA buffer containing phosphatase and protease inhibitor cocktail. Protein concentration was estimated by using the BCA kit. Equal amounts of protein were resolved by SDS-PAGE and transferred to a PVDF membrane [19, 25]. Membranes were blocked with 5% nonfat dry milk or 5% BSA followed by incubation with appropriate dilutions (1:1000) of primary antibodies overnight at 4°C and subsequently incubated with a 1:5000 dilution of horseradish peroxidase-conjugated secondary antibodies for 1 hour at room temperature. Immunoreactivity was detected by enhanced chemiluminescence solution (ImmobilonTM western, Millipore, USA) and scanned by the gel documentation system (Bio-Rad chemidoc XRS plus). To detect CXCR4 in western blot, we followed separate sample preparation as per manufacturer’s instruction.

### Apoptosis antibody array analysis

Apoptosis array was performed by using Proteome Profiler Human Apoptosis Array Kit (ARY009) from R&D Systems following manufacturer’s instructions. Detailed assay procedure was followed as described before [22]. The images were captured by the gel documentation system (Bio-Rad chemidoc XRS plus), while ImageJ software (NIH) was used for analysis. Plotly software was used for heatmap generation (Montreal, Canada).

### Real-time PCR

Total RNA was isolated from the cultured cells and tissues using the standard procedure of the RNeasy Mini Kit (Qiagen, cat no.74104). The concentration and purity of the RNA samples was determined using nanodrop. Total RNA (5 μg) of each sample was reverse-transcribed (RT) with random hexamer according to the manufacturer’s protocol (Verso cDNA synthesis kit). The final cDNA was diluted with nuclease-free water (1:3), 1μl of this having a concentration of 25ng/μl was used for each reaction in real-time PCR. Real-time PCR was carried out using an ABI StepOnePlus Real-Time PCR System (Applied Biosystems). TaqMan gene expression assay from Thermo Fisher Scientific was used for CXCR4 (Assay ID: Hs00607978_s1), CXCR7 (Assay ID: Hs00664172_s1), and DR5 (Assay ID: Hs00366278_m1) gene amplification. Reactions for each sample were performed in triplicate. GAPDH or 18s amplification was used as the housekeeping gene. A gene expression score was calculated by taking two raised to the difference in Ct between the housekeeping gene and the gene of interest (2 ΔCt). For amplifying YY1, p53, Sp1, we performed SYBR Green based RT-PCR following manufacturer’s instructions.

### Generation of stable cell lines

CXCR4 (cat#CXCR400000), CXCR7 (cat#CXCR700000), or empty vector pcDNA 3.1 (cat# V790-20) were procured from cDNA Resource Center. For stable overexpression of chemokine receptors CXCR4 and CXCR7, MCF7 cells were plated and transfected with either overexpression plasmids or empty vector individually by using Lipofectamine LTX as transfection reagent (Invitrogen). After 48 hours of transfection, cells were cultured in the presence of suboptimal dose (600µg/ml) of Geneticin (G418) for 15 days with refreshing the medium at every 3rd day for the selection of vector containing cells. The expression level for CXCR4 and CXCR7 was evaluated through flow cytometry.

3rd generation lentiviral vector pUltra-Chili-Luc (addgene no. 48688) with the bi-cistronic expression of tdTomato and Luciferase was used to make HT29 cells fluorescent. Lentiviral particles were generated in HEK-293T cells. Transduction was carried out in the presence of Polybrene (8μg/ml). A population of transduced cells (HT29-Chili-Luc) was identified by chili red expression and sorted by flow cytometry.

CXCR4 shRNA Sequence: 5’CCGGTCCTGTCCTGCTATTGCATTACTCGAGTAATGCAATAGCAGGACAGGATTTTTG 3’ was cloned into the 3rd generation transfer plasmid pLKO.1 TRC cloning vector (Addgene cat no. 10878) between unique AgeI and EcoRI sites downstream of the U6 promoter. HEK-293T cell line was used for the generation of lentiviral particles using the transfection reagent Lipofectamine 2000. The media containing the viral particles was supplemented with Polybrene (8μg/ml) for the transduction purpose. Cells were subjected to puromycin selection after 48 hours of transduction, and the knockdown profile of CXCR4 was quantified after six days of selection via Flow cytometry.

### Chromatin Immunoprecipitation (ChIP)

ChIP assay was conducted by using the ChIP assay kit (Cell Signaling Technology) following the manufacturer’s protocol. In brief, cells at 80% confluence were fixed with formaldehyde (1% final concentration directly to the culture media) for 10 minutes. Cells were then centrifuged, followed by lysis in 200μl of membrane extraction buffer containing protease inhibitor cocktail. The cell lysates were digested with MNase for 30 minutes at 37°C to get chromatin fragments followed by sonication (with 20 second on/20 second off 3 Sonication cycles at 50% amplitude) to generate 100-500 bp long DNA fragments. After centrifugation, clear supernatant was diluted (100:400) in 1X ChIP buffer with protease inhibitor cocktail followed by keeping 10% of input control apart and incubated with primary antibody or respective normal IgG antibody overnight at 4°C on a rotor. The next day, IP reactions were incubated for 2 hours in ChIP-Grade Protein G Magnetic Beads, followed by precipitation of beads and sequential washing with a low and high salt solution. Then elution of chromatin from Antibody/Protein G Magnetic beads and reversal of crosslinks by using heat was carried out. DNA was purified by using spin columns, and SYBR Green real-time PCR was conducted. Primer sequences used for the ChIP experiment for different genes are as follows: Human YY1 on DR5 activator site, Forward: 5’-CTGCCCTGACTCTCTGAAGT-3’, Reverse: 5’-GCAGACACCCCATCCCTAG-3’, Human YY1 on DR5 repressor site, Forward: 5’-TGGTTCCACACATCCCTGAA-3’, Reverse: 5’-CGCAAGCAGAAAAGGAGGTC-3’, Human p53 on DR5, Forward: 5’-CTCGAGGTCCTGCTGTTGGTGAGT-3’, Reverse: 5’-GAGCTCGGGAATTTACACCAAGTGGAG-3’

### *In-vivo* studies in xenograft tumor models

All animal studies were conducted by following standard principles and procedures approved by the Institutional Animal Ethics Committee (IAEC) of CSIR-Central Drug Research Institute. Following our well established colon cancer xenograft models as described earlier [26], 2×10^6^ cells (HT-29, DLD1, and HCT-116 control or CXCR4 KD) in 100μl PBS were subcutaneously inoculated into the flanks of the left and right hind leg respectively of each 4-6 weeks old nude Crl: CD1-Foxn1^nu^ mice. Throughout the study, the tumor was measured with an electronic digital caliper at a regular interval, and the tumor volume was calculated using standard formula V = Π / 6 × a2 × b, where ‘a’ is the short and ‘b’ is the long tumor axis. At the end of the experiment, mice were sacrificed, and subcutaneous tumors were dissected for further studies.

Parts of harvested tumors were minced into small pieces with sterile forceps in the laminar hood. These pieces of tumors are digested with collagenase, and the cells were passed through a cell strainer of pore size 70 micron to get single cells. Isolated single cells from tumor were cultured under puromycin selection for further experiments.

### Analysis of GDC TCGA Breast Cancer (BRCA) dataset

Illumina HiSeq mRNA data of Breast Cancer Cell Lines (Heiser 2012) and patients with breast cancer, GDC TCGA Breast Cancer (BRCA), was downloaded from the TCGA portal for CXCR4 and *DR5* genes using https://xena.ucsc.edu/ browser [27]. Log2 (fpkm-uq+1) values for CXCR4 and DR5 were converted into fold changes and compared to identify the association between CXCR4 and *DR5* genes. Online software Heatmapper was used for the heatmap generation, and the clustering method used was Average Linkage, whereas the Euclidean distance measurement method was considered.

### Immunofluorescent staining and confocal microscopy

Tumor samples resected from breast cancer patients (obtained from the surgical pathology at the SGPGI, Lucknow after having due Institutional ethical clearance) were fixed in 4% paraformaldehyde for 48 hours and embedded in paraffin wax. The staining of sections was performed as per the manufacturer’s recommendations (Vector Laboratories). In brief, tissue sections were de-paraffinized and rehydrated in graded alcohol. Antigen retrieval was performed by immersing slides in 10 mM sodium citrate buffer (pH 6) and boiled in a high-power microwave oven for 25 minutes. Tissue sections were then washed with PBS followed by 25-minute incubation in bloxall to neutralize endogenous peroxide activity and subsequently incubated with primary monoclonal antibodies against CXCR4 (1:800) and DR5 (1:400) in 2% BSA overnight at 4°C. The next day, the appropriate fluorochrome-conjugated secondary antibody was added and incubated for one hour. The tissue sections were then mounted by using antifade (GIBCO). Stained sections were observed under a confocal microscope (Zeiss Meta 510 LSM; Carl Zeiss). It is necessary to mention that detecting DR5 in tissue sections is challenging due to poor availability of compatible DR5 antibodies. We tried multiple antibodies for DR5 staining in tumor tissue sections and the best one is represented in the figure.

### Statistics

Most of the *in vitro* experiments are representative of at least three independent experiments. Student’s t-test and two-tailed distributions were used to calculate the statistical significance of *in vitro* and *in vivo* experiments. These analyses were done with GraphPad Prism. Results were considered statistically significant when p-values ≤ 0.05 between groups.

## Results

### CXCR4 but not CXCR7 regulates paclitaxel resistance in cancer

CXCL12 mediates its signals via CXCR4 and CXCR7 [11, 16, 28], and these signaling axes have shown to be hyperactivated in cancer with poor clinical outcome [29, 30]. Here, we sought to determine the impact of these chemokine receptors (CXCR4 and CXCR7) on chemotherapy resistance. For our gain of function studies, we selected the MCF-7 cell as it showed very negligible surface expression of both CXCR4 and CXCR7. After confirming stable CXCR4 and CXCR7 overexpression in MCF-7 cells compared to vector control (Figure 1a), we exposed cells with FDA approved anticancer drugs like Doxorubicin (250 nM), Paclitaxel (25 nM), Cisplatin (2.5 μM), and 5-Fluorouracil (25 μM) and assessed cytotoxic effects of these drugs by SRB assay. As observed in Figure 1b, selectively CXCR4 overexpressed MCF-7 cells significantly (p<0.05), resisted paclitaxel-induced cell death compared to control cells. Further, CXCR4 overexpressed cells show some degree of resistance against Doxorubicin and 5-FU, whereas similar kind of response was missing in the case of CXCR7 overexpressing cells. Next, to validate the impact of CXCR4 or CXCR7 high expression on the drug-resistant property of the cancer cells, we exploited a unique approach where we stained control MCF7 cells with DiO and CXCR4 or CXCR7 overexpressing cells with DiL respectively. DiO (green dye) and DiL (red dye) are lipophilic fluorescent dyes that get integrated into the cell membrane and transfer to the daughter cells during cell division; therefore, they offer to trace stained cells over several divisions without requiring any permeabilization or perturbing cellular function [31]. We mixed and seeded an equal number of DiO stained control cells with DiL stained CXCR4 or CXCR7 overexpressing cells. These stained cells were treated with vehicle control (DMSO) or paclitaxel (10 nmol) for five days and then monitored under either fluorescence microscopy (Figure 1c) or flow cytometry (Figure 1d). As shown in Figure 1c, green arrows indicate the DiO stained dead control cells, and the red arrows specify the DiL stained CXCR4 or CXCR7 overexpressing dead cells. The fewer number of dead CXCR4 overexpressing cells as compared to others indicate that CXCR4 contributes markedly to paclitaxel resistance. In our quantitative flow cytometry analysis, we observed that CXCR4 overexpressing MCF7 cells (DiL stained) were higher in number than control (DiO stained) cells, whereas there was no substantial change in the number of CXCR7 overexpressing cells as compared to control cells (Figure 1d). Next, we performed paclitaxel dose-response (50, 25, 12.5 nmol) study against control and CXCR4 overexpressed cells, and observed gain of CXCR4 function pose significant resistance to cell death for all three doses tested (Figure 1e). Therefore, these extensive experimental validations demonstrate that selective CXCR4 gain of function but not CXCR7 overexpression result in chemotherapy, especially paclitaxel resistance in cancer.

**Figure 1:**
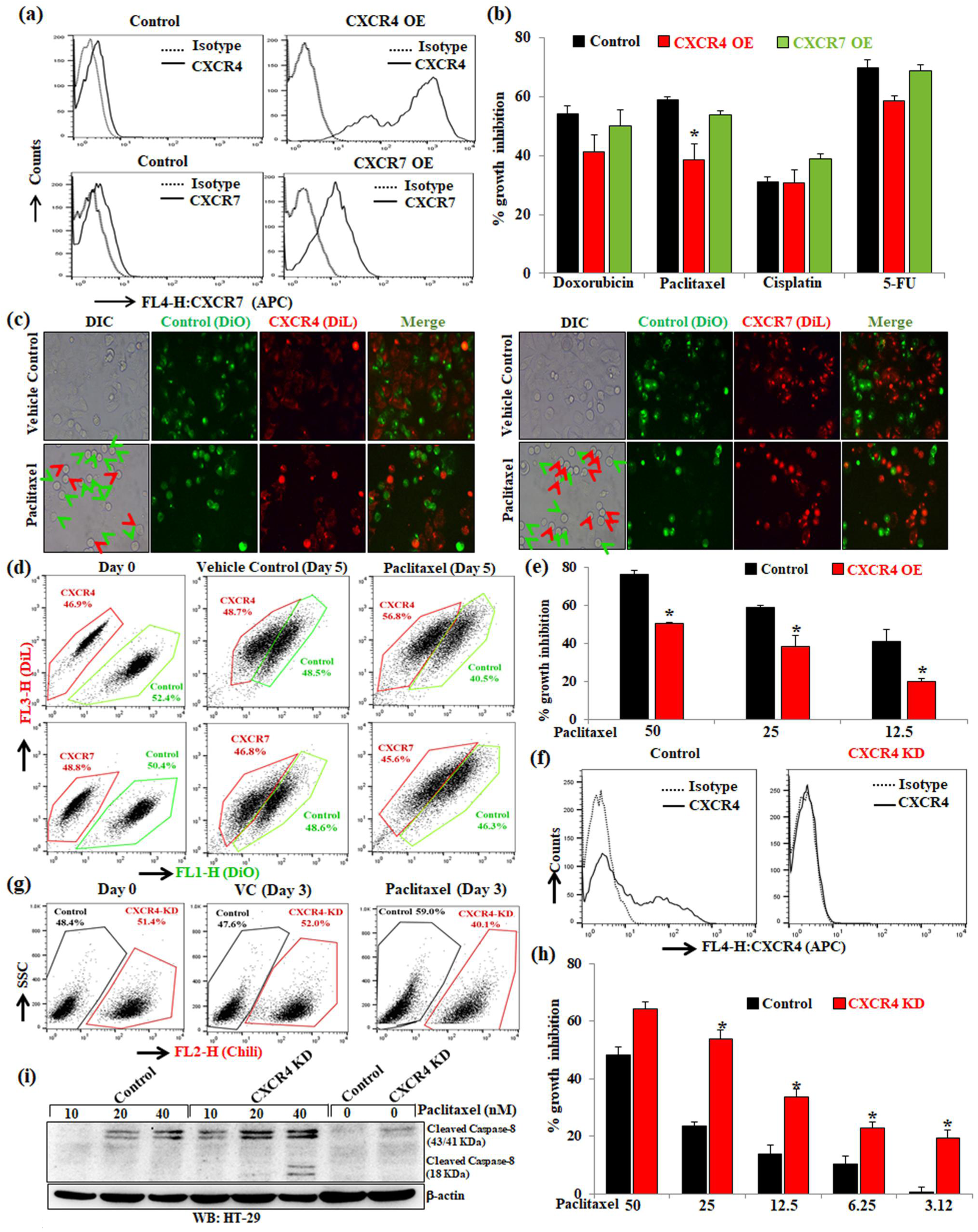
CXCR4 regulates paclitaxel resistance in cancer. (a) MCF7 cells were transfected with either empty vector pcDNA3.1 or vector containing the gene for the overexpression of either chemokine receptor CXCR4 or CXCR7 and made stable. These stable cell lines were stained with either APC-conjugated anti-human CXCR4 (CD184) or anti-human CXCR7 antibodies along with their respective isotype control antibodies and analyzed by flow cytometry. The surface expression levels of CXCR4 and CXCR7 are represented in histogram overlays. (b) CXCR4 or CXCR7 overexpressing and control MCF7 cells were treated with Doxorubicin (250 nmol/L), Paclitaxel (25 nmol/L), Cisplatin (2.5 μmol/L) or 5-Fluorouracil (25 μmol/L) for 72 hours and cytotoxicity was measured by SRB assay as described in materials and methods. Percent cell growth inhibition was tabulated. *Columns*, an average of triplicate readings of samples; *error bars*, ± SD. \**p*<0.05, compared with control cells. (c) DiL (red) stained CXCR4 or CXCR7 overexpressing MCF7 cells and DiO (green) stained control MCF7 cells were mixed in equal numbers, seeded in 6-well plate and treated with vehicle or Paclitaxel (10 nmol/L) for 3 days and then analyzed via fluorescence microscopy; red arrows indicate DiL stained dead CXCR4 or CXCR7 overexpressing cells, while green arrows indicate DiO stained dead control cells. Photomicrographs are representative of three independent experiments. (d) DiL stained CXCR4 overexpressing or CXCR7 overexpressing MCF7 cells were equally mixed with DiO stained control MCF7 cells and a small aliquot of mixture was acquired as Day 0 reading by FACS. The rest of the cells were treated either with vehicle or Paclitaxel (10 nmol/L) for 5 days and subsequently analyzed by FACS. Data are representative of at least three independent experiments. (e) CXCR4 overexpressing and control MCF7 cells were treated with different concentrations of paclitaxel (50, 25, and 12.5 nmol/L) for 48 hours and cytotoxicity was evaluated by SRB assay. Percent cell growth inhibition was tabulated. *Columns*, an average of triplicate readings of samples; *error bars*, ± SD.\**p*<0.05, compared to control cells treated with respective doses of paclitaxel. (f) Chili tagged HT-29 cells were made stable for the knockdown of CXCR4 via shRNA mediated lentiviral transduction; scramble shRNA transduced stable HT-29 cells were used as control. CXCR4 knockdown and control HT-29 cells were stained either with APC-conjugated anti-human CXCR4 (CD184) antibody or with appropriate isotype control antibody and analyzed by FACS. Histogram overlays represent the cell surface expression of CXCR4. (g) Control and CXCR4 knockdown Chili tagged HT29 cells were mixed equally and analyzed by FACS either at Day 0 or after three days of vehicle/paclitaxel (20nmol/L) treatments. (h) CXCR4 knockdown and control HT-29 cells were treated with different concentrations of Paclitaxel (50, 25, 12.5, 6.25, and 3.12 nmol/L) for 48 hours and cytotoxicity was measured by SRB assay. Percent cell growth inhibition was tabulated. *Columns*, an average of triplicate readings of samples; *error bars*, ± SD.\**p*<0.05 compared to control cells. (i) Western blot analysis of cleaved Caspase-8 in the lysate of 24 hours post vehicle or Paclitaxel treated (10, 20, and 40 nmol/L) control and CXCR4 knockdown HT-29 cells; β-Actin was used as the protein loading control. Results shown from (a) to (i) sections are representative of at least three independent experiments.

Now, we wanted to test the impact of the loss of function of CXCR4 on paclitaxel-induced cytotoxicity. Here, we select HT-29 cells for our knockdown studies as it has been shown to express CXCR4 at the basal level robustly. HT-29 cells were made stable for CXCR4 knockdown via shRNA mediated lentiviral transduction. The knockdown cells were made stable for the expression of Chili-Luc via lentiviral transduction. To check if CXCR4 loss of function sensitizes the cells to paclitaxel, equal numbers of HT-29 Chili-Luc CXCR4-knockdown cells (51.4%) and healthy control cells (48.4%) were mixed and seeded in 6-well plates. After paclitaxel treatment for three days, Chili-Luc HT-29 (CXCR4 knockdown) cells were substantially low in number (40.1%) as compared to control cells (59.0%) (Figure 1g). Next, to test the cell growth inhibitory effect of paclitaxel against CXCR4 knockdown HT-29 cells, we performed standard SRB assay and observed that paclitaxel treatment significantly (p<0.05) induces cytotoxicity in CXCR4 knockdown HT-29 cells as compared to control cells (Figure 1h). These data together suggest that the loss of CXCR4 sensitized the cells to paclitaxel treatment. To understand whether the paclitaxel-induced sensitization of CXCR4 knockdown cells is due to apoptotic cell death or not, we performed immunoblotting for Caspase-8 cleavage. It was observed that there is an induction of cleaved Caspase-8 expression in CXCR4 knockdown HT-29 cells as compared to control cells, which were further augmented upon different concentrations of paclitaxel treatment, indicating CXCR4 loss of function are priming cells for paclitaxel-induced apoptosis (Figure 1i).

### CXCR4 inversely regulates expression and function of DR5

Reciprocal regulation of chemotherapy (paclitaxel) resistance with CXCR4 expression prompted us to understand further CXCR4 regulated priming of cell death mechanisms. To explore the molecular mechanism of CXCR4 mediated therapeutic resistance, we made use of an apoptosis antibody array (ARY009, R&D Systems). This array platform is unique and an unbiased approach to study the simultaneous expression of 35 apoptosis-related proteins spanning both intrinsic and extrinsic pathways. Surprisingly, in our array hybridized with an equal quantity of proteins from CXCR4 overexpressing and control MCF7 cells, we found very selective marked reduction of expression of pro-apoptotic protein DR5 in CXCR4 overexpressed cells as compared to control cells, whereas, changes in other apoptotic proteins were found to be minimal (Figure. 2a-2b). To validate our apoptosis array results, we performed individual western blot and flow cytometry for CXCR4 overexpressed and knockdown cells along with their respective controls. As shown in Figure 2c-2f, CXCR4 inversely regulates total and surface protein expression of DR5 compared with their respective control cells. To examine the functional upregulation of DR5 under the CXCR4 knockdown condition, we determine the cytotoxic effect of TRAIL (DR5 ligand) on control and CXCR4 knockdown cells. We mixed almost equal quantity of HT-29 CXCR4 knockdown Chili-Luc cells (51.8%) and control healthy cells (47.8%) and seeded them in 6-well plates. The cells were treated with vehicle control or recombinant human TRAIL (50 ng/mL) for three days and were quantitatively analyzed through flow cytometry. As shown in Figure 2g, CXCR4 knockdown Chili-Luc cells were comparatively less in number (36.5%) as compared to control cells (63.1%), suggesting that CXCR4 knockdown resulted in induction of functional DR5 on the cancer cell surface. Next, we determined the dose-response of rhTRAIL against control and CXCR4 KD cells and observed a significant increase in cell death in CXCR4 KD cells compared to control (Figure 2h). To test TRAIL-mediated execution of apoptosis in cancer cells, we performed immunoblotting for cleaved Caspase-8 expression and noticed that rhTRAIL was found to be robustly effective in inducing cleavage of Caspase-8 in CXCR4 KD cells compared to their respective controls. Altogether, our intricate experimental planning demonstrates that CXCR4 inversely alters the expression of DR5, and loss of its function results in TRAIL-mediated sensitization of cancer cells.

**Figure 2.**
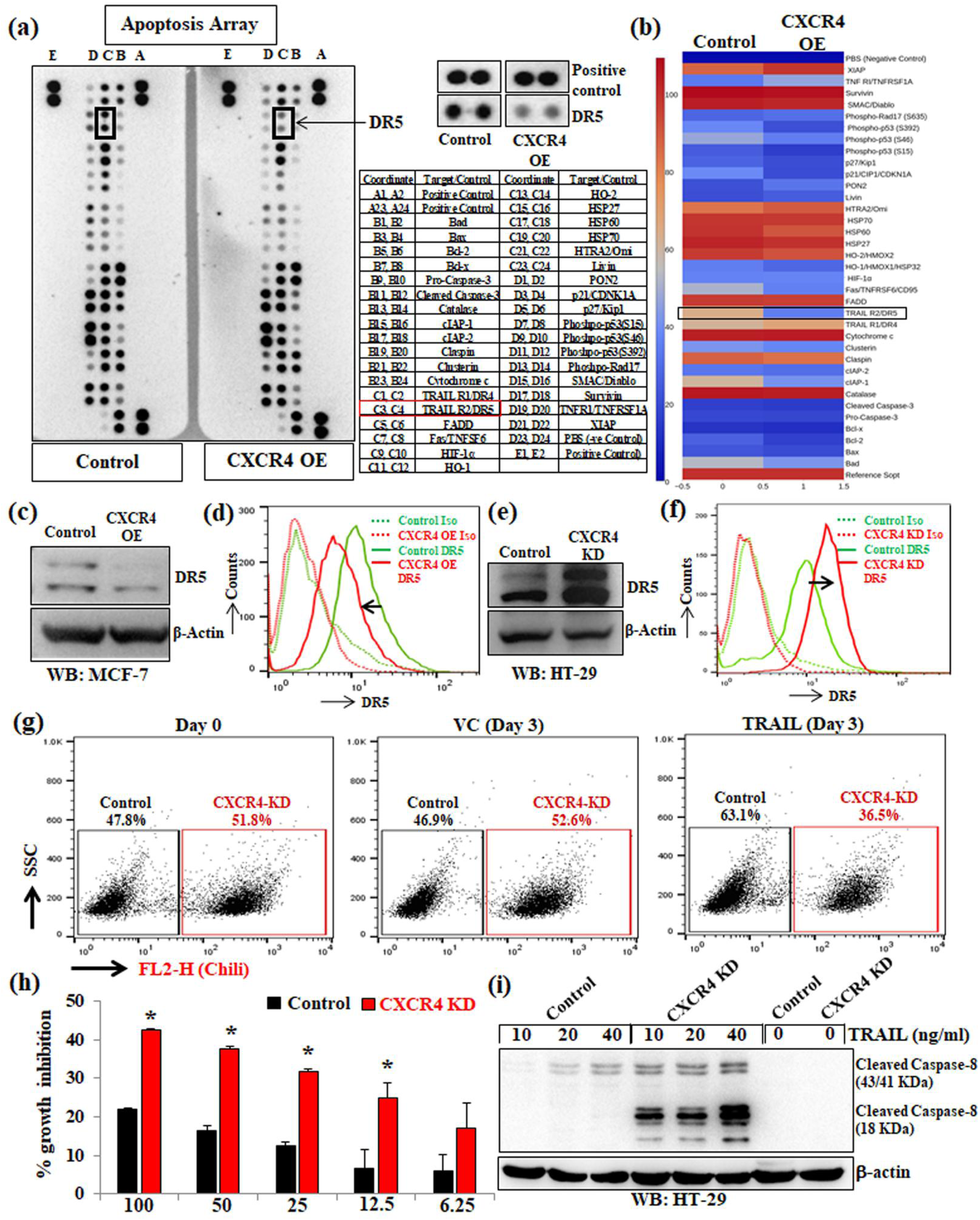
CXCR4 inversely regulates the expression and function of DR5. CXCR4 overexpressing or control stable MCF7 cells were harvested for protein extraction and analyzed for the expression of apoptotic genes by utilizing proteome profiler apoptosis array or individual western blot analysis. (a) Chemiluminescent image of the expression of 35 apoptosis related genes with positive and negative controls in duplicates for control and CXCR4 overexpressed cells were shown (left panel). The enlarged images of selected apoptotic protein (DR5) found to be markedly altered in the proteome profiler array (middle upper) and spot coordinates (middle lower) were shown. (b) Heatmaps depicting differentially regulated proteins in CXCR4 overexpressing and control stable MCF7 cells. (c) Immunoblot analysis of DR5 protein in CXCR4 overexpressing or control MCF7 cells; β-Actin was used as an internal protein loading control. (d) CXCR4 overexpression and control MCF7 cells were either stained with PE-conjugated anti-human DR5 or PE tagged isotype (IgG) control antibodies and cell surface expression of DR5 was analyzed by histogram overlays using FACS. (e) Immunoblot analysis of DR5 protein in CXCR4 knockdown or control HT-29 cells; β-Actin was used as an internal protein loading control. (f) CXCR4 knockdown and control HT-29 stable cells were stained either with PE-conjugated anti-human DR5 or PE tagged isotype control antibodies. Cell surface expression of CXCR4 was analyzed by histogram overlays using FACS. (g) Chili tagged CXCR4 knockdown and control HT-29 stable cells were mixed equally and subjected to FACS analysis at Day 0 and 3 days after recombinant human TRAIL (50 ng/mL) treatment. (h) CXCR4 knockdown and control stable cells were treated with different concentrations of TRAIL (100, 50, 25, 12.5, and 6.25 ng/mL) for 48 hours and cytotoxicity was measured by SRB assay. Percent cell growth inhibition was tabulated. *Columns*, an average of triplicate readings of samples; *error bars*, ± S.D. **p < 0.01; #, p < 0.05, compared to TRAIL-treated control cells. (i) Immunoblot analysis of cleaved Caspase-8 in 24 hours post vehicle or TRAIL-treated (10, 20, and 40 ng/mL) control and CXCR4 knockdown HT-29 cell lysates; β-Actin was used as the protein loading control. Results shown from (a) to (i) sections are representative of at least three independent experiments.

### CXCR4 mediated DR5 downregulation is dependent on the recruitment of transcription factors p53 and YY1 to its promoter

To ascertain that the alteration of DR5 expression is at the transcriptional level also, mRNA expression was assessed by real-time PCR in control, and CXCR4 overexpressed/knockdown cells. As shown in Figures 3a and 3b, mRNA expression of the *DR5* gene (TNFRSF10B) was found to be significantly (p<0.05) lower in CXCR4 overexpressing cells as compared to control cells and vice-versa in knockdown cells as compared to their respective control. To identify the transcription factors contributing towards the transcriptional regulation of DR5 under CXCR4 expression, we went through the literature and found that p53, YY1, and Sp1 are major transcriptional regulators of DR5 as they have specific binding sites on the promoter region of *DR5* gene. p53 and Sp1 are the positive regulators, while YY1 is a negative regulator of *DR5* gene transcription [32-34]. In order to check the status of these transcription factors in our cells following CXCR4 overexpression, we performed immunoblotting and observed that p53 was found to be downregulated. In contrast, the expression of YY1 goes up under the influence of CXCR4 overexpression as compared to the controls (Figure 3c). Compared to control, significant (p<0.05) downregulation of p53 mRNA and upregulation of YY1 mRNA was observed after CXCR4 overexpression as determined by real-time qPCR analysis (Figure 3d). These data indicate that the DR5 regulating transcription factors, p53 and YY1, are themselves regulated at the transcriptional level in response to the overexpression of CXCR4 in cancer cells. Next, to analyze the binding of these DR5 regulating transcription factors on the *DR5* gene promoter region in CXCR4 overexpressing and control cells, we performed promoter analysis for putative binding sites of YY1 and p53 on DR5 promoter by using the MatInspector tool from Genomatix software. We designed the primers for these protein binding sites on the DR5 promoter. By using the software, we found one activator binding site for p53, whereas one repressor binding site and one activator binding site of YY1 on the DR5 promoter region (Figure 3e). So, we performed Chromatin Immunoprecipitation (ChIP) assay of CXCR4 overexpressing and control MCF7 cells and pull down the p53 (or YY1) associated chromatin by using human anti-p53 (or anti-YY1) antibody. Co-precipitated DNA was eluted from the pulled-down chromatin, and real-time qPCR was performed for amplification of associated pulled-down DNA. We found the decreased binding of p53 on the DR5 promoter and increased recruitment of YY1 on the repressor site; while decreased binding of YY1 on the activator site of the *DR5* gene promoter in CXCR4 overexpressing cells as compared to control cells (Figure 3f). Thus, from the overall results of immunoblot, real-time PCR and ChIP assays, it is clear that the downregulation of DR5 in CXCR4 overexpressing cells is mediated by the decreased recruitment of p53 on its activator site and increased recruitment of YY1 on its repressor site of the DR5 promoter region.

**Figure 3.**
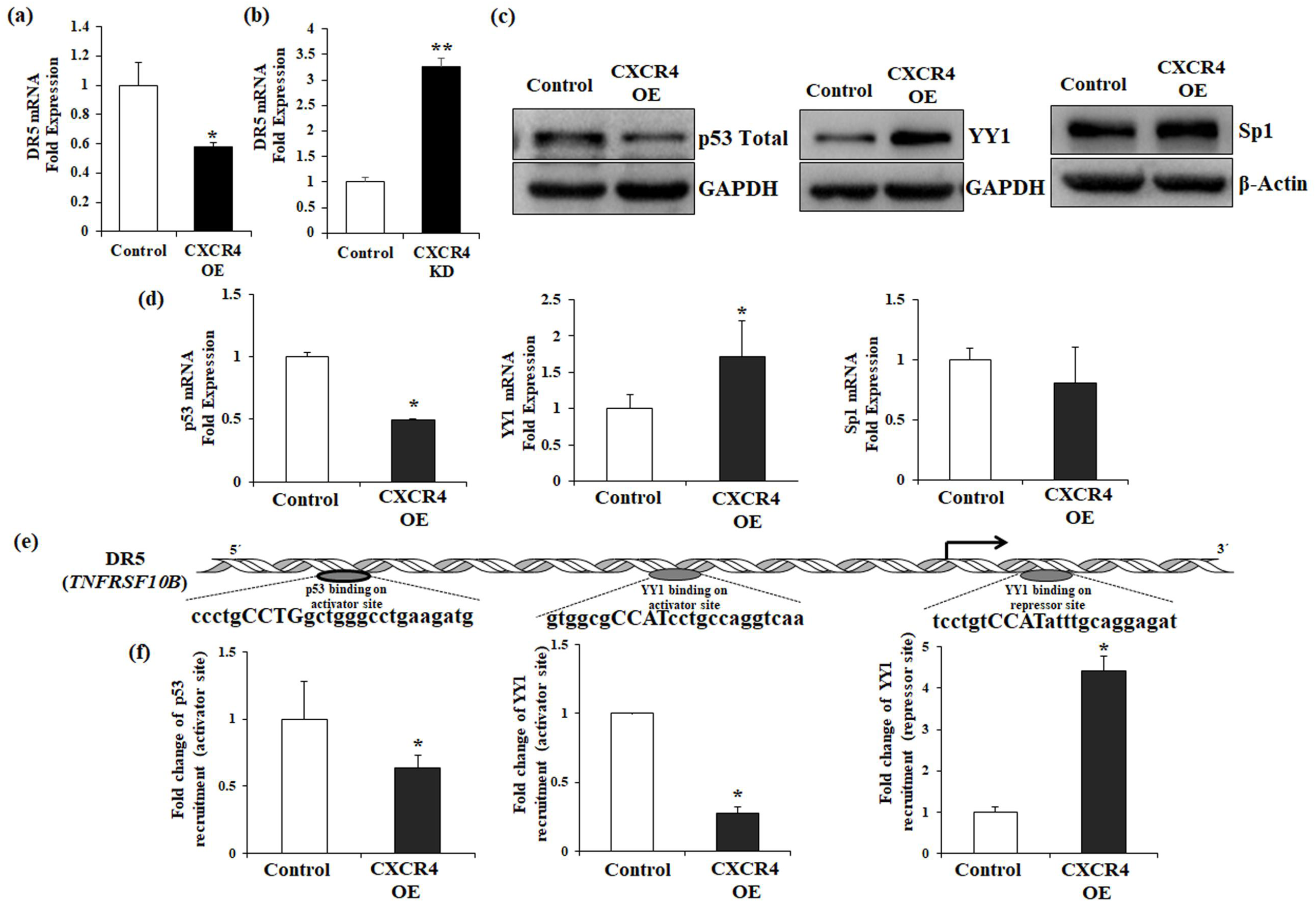
CXCR4 regulates DR5 transcription by differentially modulating the recruitment of transcription factors p53 and YY1 at the promoter site of DR5. (a-b) Total RNA was isolated from CXCR4 overexpressing (MCF-7) and knockdown (HT-29) stable cells along with their respective controls and reverse transcribed. Fold change in DR5 mRNA expression was measured by real-time PCR as described in Materials and Methods. Data are representative of three independent experiments, resulting from duplicate readings of two different samples; Columns, average value of DR5 mRNA expression; bars, ±SD. *, p < 0.05, compared with respective control. (c) Western blot analysis of p53, YY1, and Sp1 in control and CXCR4 overexpressing MCF-7 cells; GAPDH or β-actin was used as protein loading control. (d) Fold change in mRNA expression of p53, YY1, and Sp1 in control and CXCR4 overexpressing stable MCF-7 cells was assessed by real-time PCR. Results are representative of three independent experiments; *Columns*, average value of p53/yy1/sp1 mRNA expression; bars, ±SD. *, p < 0.05, compared with respective control. (e) Diagrammatic representation for the p53 binding on activator site as well as YY1 binding on activator and repressor sites of the *DR5* gene promoter region. (f) ChIP assay for the analysis of YY1 and p53 recruitment on the *DR5* gene promoter in CXCR4 overexpressing and control stable MCF7 cells followed by real-time PCR. Fold change in p53 and yy1 recruitment on activator site (left and middle) as well as yy1 recruitment on repressor site (right) of the *DR5* gene promoter were assessed in control and CXCR4 overexpressing MCF-7 cells. Results are representative of at least two independent experiments; *Columns*, an average of duplicate readings of samples; *error bars*, ± S.D. \**p* < 0.05 versus control MCF7 cells.

### CXCL12-CXCR4 mediated signaling pathway is not responsible for DR5 regulation and paclitaxel sensitization

Reverse regulation of cancer cell apoptosis and functional expression of pro-apoptotic protein DR5 under CXCR4 gain and loss of function encouraged us to test the contribution of CXCR4 ligand CXCL12 mediated signaling axis for delivering the above phenotype. First, we attempted to quantify CXCL12 in the culture supernatant of control and CXCR4 overexpressed MCF-7 cells, however, to our surprise, we were unable to detect any CXCL12 in the medium (data not shown). This unexpected result prompted us to carry out further experiments to determine the contribution of CXCL12-CXCR4 signaling axis in pro-tumorigenic functions of CXCR4 observed earlier. To block CXCR4 mediated signals, we used FDA approved CXCR4 antagonist AMD3100 [35, 36]. We treated our control and CXCR4 overexpressed MCF-7 cells either with AMD3100 or CXCL12 and sought to observe the change in suppressed DR5 expression. Unexpectedly, compared to control, we did not observe any rescue (in case of AMD3100 treatment) or further suppression (CXCL12 treatment) of DR5 expression following respective treatments (Figure 4a). To further validate our previous observations, we selected the HT-29 cell, which has been shown to highly express CXCR4 on its cell surface. In order to mimic the phenotype of our CXCR4 knockdown experiments, HT29 cells were treated with AMD3100 and paclitaxel alone and in combination to assess any synergistic cytotoxic effect of paclitaxel under CXCL12-CXCR4 inhibited condition. However, to our great surprise, we did not find any significant synergistic cytotoxic effect in combination treatment (AMD3100 plus paclitaxel) compared to individual treatments (Figure 4a). Two entirely unexpected observations forced us to test the efficacy of AMD3100 and CXCL12 in our system. As ERK phosphorylation is a classical downstream signature of the CXCR4-CXCL12 axis, we examined the effect of CXCL12 and AMD3100 on the regulation of phosphorylation of ERK in HT29 cells. As expected, CXCL12 mediated ERK phosphorylation was found to be markedly inhibited by AMD3100 treatment, strongly advocating the fact that both CXCL12 and AMD3100 are fine and functional in our system and confirmed the accuracy of our previous unexpected observations (Figure 4c). To further rule out the possible involvement of CXCL12-CXCR4 signaling in modulating DR5 expression, we sorted CXCR4^+^ and CXCR4^−^ cells by FACS from HT29 cells, allowed them to grow for five days and analyzed the expression of DR5 in Day 0 and Day 5. As shown in Figure 4d, upper and lower panels, sorted cells maintained their CXCR4^+^ and CXCR4^−^ status, however, there was no substantial difference in the expression of DR5 in the beginning and fifth day of culture. To further validate our unique observations in a more universal manner, we examined the surface expression of CXCR4 in several different cancer cell lines and selected some like DLD-1, HCT-116, A-549, and MDA-MB-468, where there is negligible or no surface expression of CXCR4 (Figure 4e), however, CXCR4 protein is present in all the cases in cytoplasm as observed by Western blot analysis. Here, it is noteworthy to mention that we were not able to develop CXCR4 and DR5 in the same blot as detecting CXCR4 protein in western blot requires a particular sample preparation condition in which DR5 cannot be detected. Most interestingly, CXCR4 knockdown in all the cases markedly induced the expression of DR5 across various cancer cell lines of human origin (Figure 4f), strongly suggesting intracellular CXCR4 protein inversely regulates the expression of DR5 in cancer. Altogether, our extensive well-connected experiments demonstrate that intracellular CXCR4 protein, but not CXCR4-CXCL12 mediated signals regulate the expression of DR5 in cancer.

**Figure 4.**
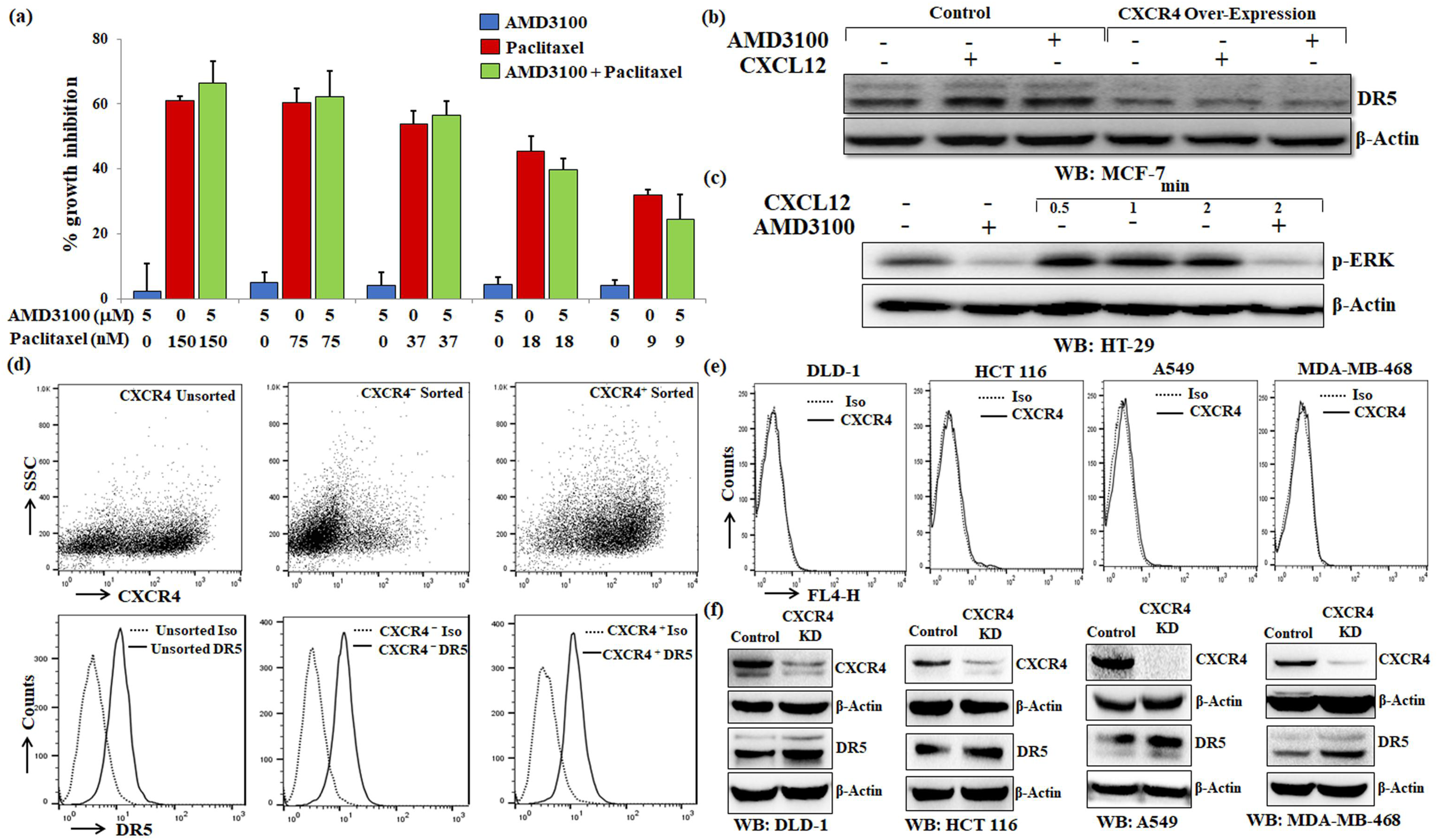
CXCR4 protein but not its ligand (CXCL12) mediated signals regulate the expression of DR5. (a) Control and CXCR4 overexpressed MCF-7 cells were treated with CXCR4 ligand CXCL12 (100ng/ml) or CXCR4 antagonist AMD3100 (5 μmol/L) for 12 hours, and subjected to Western blot analysis for DR5 and β-actin. (b) HT-29 cells were treated with either different concentrations of paclitaxel (150, 75, 37.5, 18 and 9 nmol/L) or AMD3100 (5 μmol/L) alone or in combinations for 48 hours and cytotoxicity was measured by SRB assay. Percent cell growth inhibition was tabulated. *Columns*, an average of triplicate readings of samples; *error bars*, ± SD. (c) HT-29 cells were either pre-treated with vehicle or AMD3100 (5 μmol/L) for 12 hours followed by treatment with CXCR4 ligand CXCL12 (100ng/ml) for different time points (0.5, 1 and 2 mins) and subjected to Western blot analysis for p-ERK and β-actin. (d) CXCR4^+^ and CXCR4^−^ HT-29 cells were flow sorted and plated. After 5 days of culture, cells were stained with either APC-conjugated CXCR4 (CD184) and PE-conjugated DR5 or their respective matched isotype control antibodies and analyzed by FACS. In upper panel, dot plots represent CXCR4 staining in unsorted, CXCR4^+^ sorted and CXCR4^−^ sorted cells. In lower panel histograms represent DR5 staining in the above-mentioned respective cells. (e) DLD-1, HCT-116, A-549, and MDA-MB-468 cells were stained with either APC-conjugated anti-human CXCR4 (CD184) or isotype control antibodies and analyzed by FACS. The cell surface expression of CXCR4 is represented in histogram overlays. (f) DLD-1, HCT-116, A-549, and MDA-MB-468 cells were made stable for CXCR4 knockdown via shRNA mediated lentiviral transduction and scramble shRNA transduced cells were used as control. Immunoblot analysis of CXCR4 and DR5 protein in control or CXCR4 knockdown cells are shown, β-Actin was used as an internal protein loading control. Results shown from (a) to (f) sections are representative of three independent experiments.

### Loss of CXCR4 protein results in compromised colon tumor growth *in vivo*

A series of earlier *in vitro* experiments across diverse human cancer types suggested CXCR4 protein, but not its ligand-mediated signals are critical for CXCR4 mediated paclitaxel resistance and CXCR4 protein inversely regulates expression of pro-apoptotic protein DR5. To validate our atypical *in vitro* observation into an *in vivo* system, we picked three different colon cancer cell lines having different CXCR4 expression patterns, such as HT-29 have robust CXCR4 surface expression and other two DLD-1 and HCT-116 having only intracellular CXCR4 expression. To investigate the contribution of CXCR4 surface expression versus intracellular expression in modulating *in vivo* tumor growth, we inoculated 2 million control or CXCR4 knockdown cells of all three (HT-29, DLD-1 and HCT-116) into the right or left hind leg of 4-6 weeks old nude Crl: CD1-Foxn1nu mice and assessed tumor progression. Remarkably, in all three xenograft animal models, irrespective of surface expression of CXCR4, loss of receptor function through CXCR4 knockdown resulted in significant inhibition of progression of tumor growth compared to their respective control cells (Figure 5a-5d and supplementary Figure 1). FACS analysis of single cells isolated from CXCR4 knockdown HT-29 xenograft tumors displays maintenance of CXCR4 knockdown status whereas, single cells from control HT-29 tumors exhibit a similar extent of basal positivity of CXCR4 surface expression (Figure 5e). CXCR4 surface expression was found to be almost nil in single cells isolated from both control and CXCR4 knockdown DLD-1 xenograft tumors (Figure 5f). However, still, CXCR4 knockdown caused robust tumor growth compromisation suggesting CXCR4 surface expression has no impact on the progression of colon tumorigenesis. To check the status of DR5 in the *in vivo* state, we performed western blot analysis in harvested tumors and observed the marked upregulation of the expression of DR5 in the CXCR4 knockdown condition compared with their respective controls (Figure 5g, 5h). Together, our xenograft experiments in the animal also suggest that CXCR4 protein, but not its ligand-mediated signals are essential in modulating *in vivo* tumor growth.

**Figure 5.**
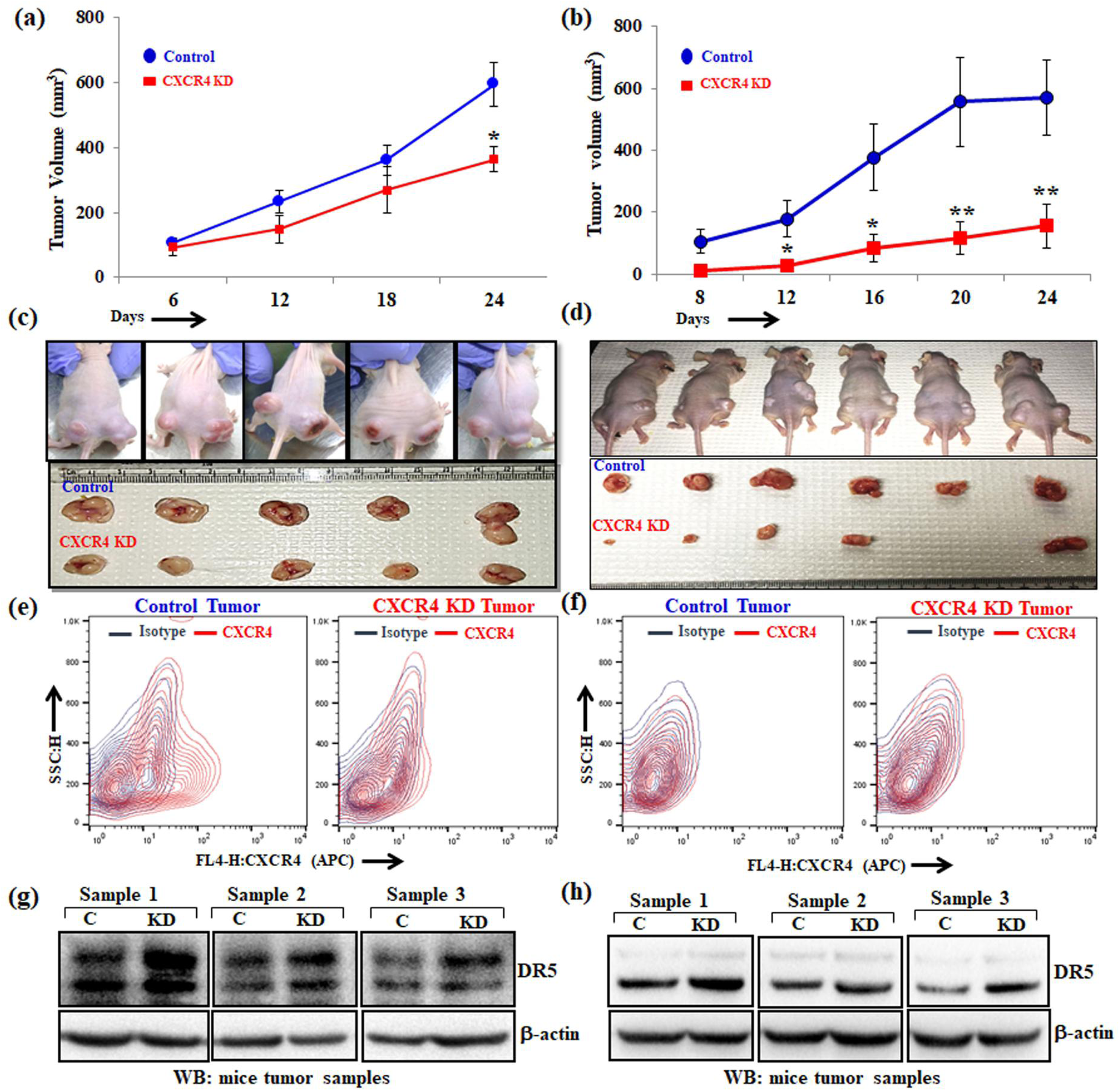
CXCR4 protein knockdown results in compromised tumor growth and DR5 overexpression *in vivo*. 2 ×10^6^ stable control (HT-29 and DLD-1) or CXCR4 knockdown (HT-29 and DLD-1) cells in 100μl PBS were injected subcutaneously in the flanks of the right or left hind leg of 4-6 weeks old Crl:CD1-Foxn1^nu^ mice respectively. Tumor volumes were measured after regular intervals by using a caliper. Growth curves for HT-29 (a) and DLD-1 (b) are shown for tumors generated from control and CXCR4 knockdown cells; points are indicative of average value of tumor volume (n=5 for HT-29, n=6 for DLD-1); bars, **±** SE.. \**p*<0.05 compared to control tumors. (c and d) Upper panels represent images of tumor bearing mice, control (right flank) and CXCR4 knockdown (left flank). Mice were sacrificed, and the respective tumors from HT29 (c) and DLD-1 (d) were harvested and shown in photographs in lower panels. Single cells from respective control and CXCR4 knockdown HT-29 (e) and DLD-1 (f) were harvested and stained with either APC-conjugated anti-human CXCR4 (CD184) or isotype control antibodies and cell surface expression of CXCR4 was analyzed by contour FACS plots. Harvested tumors generated from control and CXCR4 knockdown cells [HT-29 (g) and DLD-1 (h)] were subjected to Western blot analysis for DR5 and β-actin.

### Expression of CXCR4 and DR5 are inversely correlated in human breast cancer samples

To understand the pathophysiological significance of our finding in context of human cancer, we exploited TCGA database to find out clinical correlation between CXCR4 and DR5, especially focusing on breast cancer as it possesses large (n=54) panel of breast cancer cell line data along with significant amount of breast cancer patient (n=1217) sample data [27]. First, we analyzed the expression profiles of CXCR4 and DR5 by using UCSC Xena (https://xena.ucsc.edu/) browser in a panel of breast cancer cell lines and found an astonishing inverse correlation between CXCR4 and DR5 expression in most of the breast cancer cell lines (Figure 6a). As panel cell line data is not available for human colon and other cancers, we further analyzed CXCR4 and DR5 expression by real-time PCR in 7 different cancer cell lines, four of which belongs to colon cancer and observed an inverse association between CXCR4 and DR5 expression in most of the solid tumor cell lines (Figure 6b). Next, we extended our similar studies covering a large number of human breast cancer tissue samples and found a marked reverse relationship between CXCR4 and DR5 expression (Figure 6c). To validate TCGA data, we next evaluated the mRNA expression of CXCR4 and DR5 in human breast tissues obtained from eight patients with breast carcinoma by real-time PCR, which demonstrates that CXCR4 expression is oppositely regulated with DR5 expression. Finally, we validated our observations at the protein level by performing immunofluorescence microscopy in our procured breast cancer samples, where we double-stained CXCR4 (red) and DR5 (green). As observed in Figure 6e, CXCR4 protein expression was found to be predominant in human breast cancer tissues as expected, whereas, DR5 expression was sparse. Interestingly, even a few cells in breast tumor tissue samples that were found to be positive for more DR5 expression are devoid of CXCR4 expression suggesting the inverse relationship between them again. Altogether, these findings indicate that our *in vitro* observations are likely of pathophysiologic importance for the development of solid tumors *in vivo*, where the inverse relationship between CXCR4 versus DR5 could be a critical indicator for predicting therapy response as well as the aggressiveness of particular cancer.

**Figure 6.**
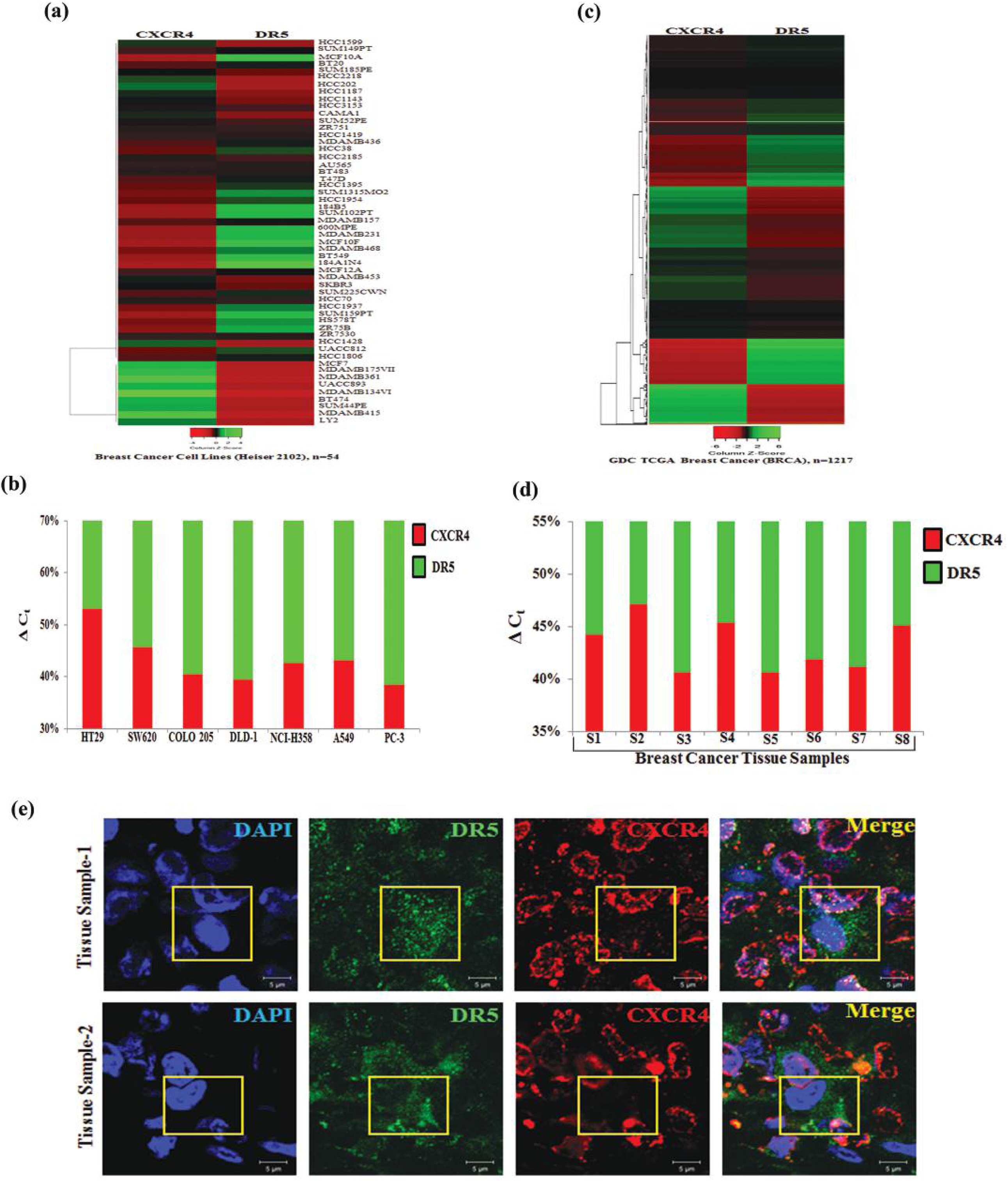
Expression of CXCR4 inversely correlates with DR5 expression in cancer cell lines, human breast cancer patient cohort and tissue samples. Heatmaps displaying CXCR4 and DR5 expression in (a) breast cancer cell lines (n=54) and in (b) GDC TCGA Breast Cancer (BRCA) patient cohort (n=1217). Shades of red and green represent expression values in fold change. Total RNA was isolated from various cancer cell lines (c) and breast cancer patient tumor tissue samples (d), reverse transcribed and real-time PCR was performed for CXCR4 and DR5 expression analysis. 18s is used as an internal control. Percentage delta C_t_ was determined for each sample from quadruplicate C_t_ value and represented in bar graph having the differential contribution of CXCR4 (red) and DR5 (green) expression. (e) Human formalin-fixed paraffin-embedded mammary tumor tissues were subjected to immuno-fluorescence staining of CXCR4 (red) and DR5 (green) proteins and analyzed by confocal microscopy. Scale bar, 5 μ Sections were viewed at 63 X magnification. Yellow boxed merged confocal photomicrograph area represents cell positive for DR5 (green) staining having minimal CXCR4 (red) staining.

## Discussion

The lineage of CXCR4 with its pro-tumorigenic functions in solid tumors is unequivocal. Earlier publications suggest that the CXCR4-CXCL12 axis is not only pivotal for modulating cancer metastasis but also responsible for executing its other tumor-promoting functions [11, 37, 38]. Despite vast preclinical evidence, CXCR4 inhibitor ‘Plerixafor’ or AMD3100 got FDA approval as a stem cell mobilizer [39], not as a cancer drug or even as a metastasis inhibitor. The discrepancy of preclinical observations and limited clinical success of CXCR4 antagonists as cancer therapy strongly advocates the involvement of some other aspects of CXCR4 biology beyond its classical CXCR4-CXCL12 signaling axis. Our data for the first time, suggests that intracellular presence of CXCR4 protein, but neither its surface expression nor CXCL12-CXCR4 mediated signals are essential in modulating therapy (paclitaxel) resistance in cancer. Our results also justify to some extent why the CXCR4 inhibitor phenotype does not match with the CXCR4 null/knock-out phenotype under different preclinical settings.

Initially, we considered both CXCR4 and CXCR7 to study the impact of these two chemokine receptors to understand therapy resistance in cancer as they share common ligand CXCL12, and are shown to be overexpressed in different solid tumors [11, 14]. However, we observed selective involvement of CXCR4 in significantly modulating paclitaxel resistance in breast and colon cancers, which is actually in corollary with vast literature documenting the pro-tumorigenic role of CXCR4. Drug resistance is one of the identifying features of Cancer stem cells (CSCs) [40-43] and CXCR4; being a bona fide CSC marker for prostate and pancreatic cancers [44, 45], evidently support its presence to promote therapy resistance. Also, studies in AML and NSCLC showed the presence of CXCR4 has a positive correlation with therapy resistance though they consider ligand-mediated signals as a responsible reason for the same [46, 47]. Our unbiased mechanistic hunt discovered that CXCR4 mediates therapy resistance via selectively regulating the pro-apoptotic candidate protein DR5 by differentially modulating YY1 recruitment at the promoter site of the DR5 gene. Differential YY1 recruitment in the promoter region for activation and repression of target genes is documented by earlier elegant studies [48, 49]. Further, CXCR4-YY1 reciprocal regulation has been well documented as a pro-tumorigenic function of CXCR4 in AML therapy resistance and osteosarcoma angiogenesis [46, 50]. CXCR4 is rarely found to be localized in the nucleus which has been shown to be linked with poor prognosis and enhanced metastasis [51, 52]. Nuclear localization of CXCR4 may be associated with transcriptional upregulation of YY1 and its differential recruitment to the promoter region of *DR5* gene. Though CXCR4 mediated DR5 transcriptional regulation is our novel finding, it has been reported in the studies that high CXCR4 expression in the cancer cells is correlated with poor prognosis and resistance against the various DNA damaging chemotherapeutic agents whose mechanism of action involve the regulation of Death receptors [53, 54].

Utilizing three different xenograft models of colon cancer cells that are either expressing surface CXCR4 (HT-29) or are null for the CXCR4 surface expression (DLD-1, HCT-116), we provided strong evidence that knockdown of CXCR4 results in reduced tumor growth irrespective of their surface expression status (Figure 5 and supplementary Figure 1). In support of our *in vivo* observations, several previous studies have demonstrated that CXCR4 knockdown cells produce smaller tumors as compared to their control counterparts [38, 55, 56]. Interestingly, at least one study indicated that the cytoplasmic expression of CXCR4 is correlated with tumor burden and the metastatic load of certain cancers [57]. Further, some reports have suggested that the *in vivo* environment gives cues to the cancer cells to transport their intracellular CXCR4 on the surface [37, 58], so to test the same, we isolated single cells from harvested *in vivo* xenograft tumors and examined the CXCR4 surface expression. However, no change was found in the surface expression of CXCR4 either in control or the knockdown cells suggesting the fact that though they gave rise to smaller tumors compared to control, there is no contribution of CXCR4-CXCL12 signaling axis in delivering this phenotype. Inverse correlation of CXCR4 and DR5 expression was observed in our human breast cancer samples, which was primarily allied with TCGA data obtained from a broad panel of breast cancer cell lines as well as data from human TCGA Breast Cancer (BRCA) cohort suggesting the clinical significance of our finding.

Overall, the study indicates that high expression of CXCR4 protein but not CXCR4 signaling restricts cellular apoptosis and promotes cell survival via downregulating the DR5 expression and thus renders cancer cells resistant to chemotherapeutic drugs. Further, intracellular CXCR4 protein contributes significantly to the tumorigenic potential of cancer cells, and future therapies should majorly focus on targeting CXCR4 protein and not only CXCR4 mediated signals. Targeting selectively CXCR4 protein in cancer cells would not be easy. However, our findings open up a huge possibility for the further discovery of CXCR4 interacting protein partners, which might have an immense role in posing its pro-tumorigenic effect and can be effectively targeted by small molecule inhibitors. Moreover, our original finding may trigger the discovery of the novel intracellular role of other chemokine receptors in the context of particular disease pathophysiology, which has been overlooked so far.

## Acknowledgments

We sincerely acknowledge the excellent technical help of Mr. A. L. Vishwakarma of SAIF for the Flow Cytometry studies; Dr. K. Singh and Dr. K. Mitra of Electron Microscopy unit for Confocal Imaging and Mr. S. Meena for providing the routine cell culture facilities. Research of all the authors’ laboratories was supported by CSIR-CDRI Institutional Fund and Fellowship grants from CSIR, DBT and UGC. D.D acknowledges grant support from DST (EMR/2016/006935) and DBT (BT/AIR0568/PACE-15/18). Institutional (CSIR-CDRI) communication number for this article is 135.

## Author Contributions

MAN and SM involved in study designing, performed experiments and wrote the draft manuscript. AS, AKS, RKA, PC, AV and KKS provided active support for carrying out various *in vitro* and *in vivo* experiments. AG and AM provided the human tissue samples. DD conceived the idea, designed experiments, analyzed data, wrote the manuscript and provided overall supervision. All authors read and approved the final manuscript.

