## Supplementary Figure 1 for "CXCR4 intracellular protein regulates drug resistance and tumorigenic potential via modulating the expression of Death Receptor 5"

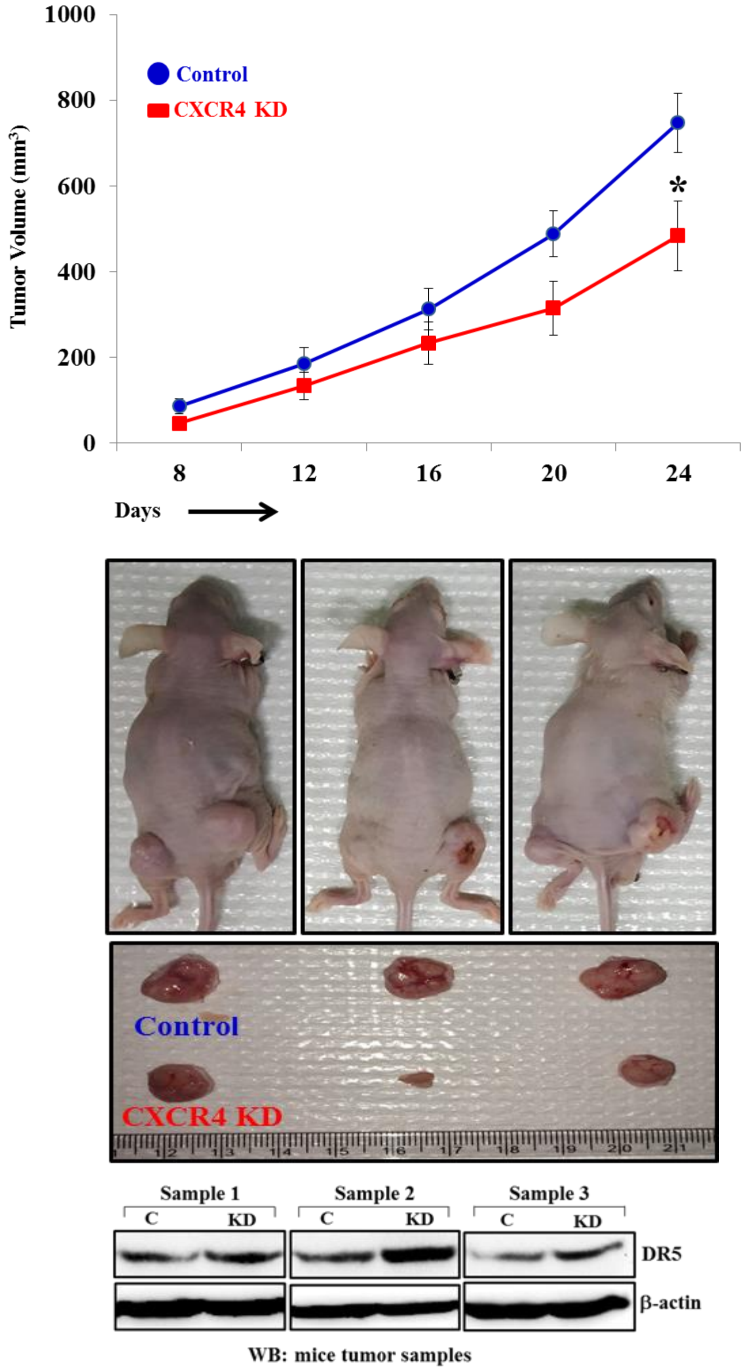


**Supplementary Figure 1: CXCR4 protein knockdown results in compromised tumor growth and DR5 overexpression *in vivo* in HCT-116 xenograft model**

2 X$10^6$ stable control or CXCR4 knockdown HCT-116 cells in 100µl PBS were injected subcutaneously in the flanks of the right or left hind leg of 4-6 weeks old Crl:CD1-Foxn1$^{nu}$

mice. Tumor volumes were measured after regular intervals by using a caliper. Tumor growth curves are shown; points are indicative of average value of tumor volume (n=10); bars**,** ± SE. *$p<0.05$ compared to control tumors. Middle panel represent images of tumor bearing mice, control (right flank) and CXCR4 knockdown (left flank). Mice were sacrificed, and the respective tumors were harvested and shown in photographs. Harvested tumors generated from control and CXCR4 knockdown cells were subjected to Western blot analysis for DR5 and β-actin (lower panel).
